# PopGenAdapt: Semi-Supervised Domain Adaptation for Genotype-to-Phenotype Prediction in Underrepresented Populations

**DOI:** 10.1101/2023.10.10.561715

**Authors:** Marçal Comajoan Cara, Daniel Mas Montserrat, Alexander G. Ioannidis

## Abstract

The lack of diversity in genomic datasets, currently skewed towards individuals of European ancestry, presents a challenge in developing inclusive biomedical models. The scarcity of such data is particularly evident in labeled datasets that include genomic data linked to electronic health records. To address this gap, this paper presents PopGenAdapt, a genotype-to-phenotype prediction model which adopts semi-supervised domain adaptation (SSDA) techniques originally proposed for computer vision. PopGenAdapt is designed to leverage the substantial labeled data available from individuals of European ancestry, as well as the limited labeled and the larger amount of unlabeled data from currently underrepresented populations. The method is evaluated in underrepresented populations from Nigeria, Sri Lanka, and Hawaii for the prediction of several disease outcomes. The results suggest a significant improvement in the performance of genotype-to-phenotype models for these populations over state-of-the-art supervised learning methods, setting SSDA as a promising strategy for creating more inclusive machine learning models in biomedical research.

Our code is available at https://github.com/AI-sandbox/PopGenAdapt.

## 1. Introduction

Genomic data has become increasingly important for biomedical research, as it can reveal insights into the causes, diagnosis, prevention, and treatment of various diseases. However, the available data is predominantly from individuals of European ancestry, despite their making up only 16% of the global population. This disproportionate representation presents one of the major challenges in developing biomedical models and studies that can effectively generalize across diverse populations, posing the risk of exacerbating existing health disparities.^1^ While widely adopted datasets such as the UK Biobank^2^ provide rich phenotypic information from electronic health records, they lack diversity (see Fig. 1). On the other hand, highly diverse datasets, such as gnomAD,^3^ lack phenotypic data, which makes them not directly usable to train supervised genotype-to-phenotype machine learning models, as phenotype labels for all the samples are required. New algorithmic solutions are needed in order to profit from all available data.

**Fig. 1.**
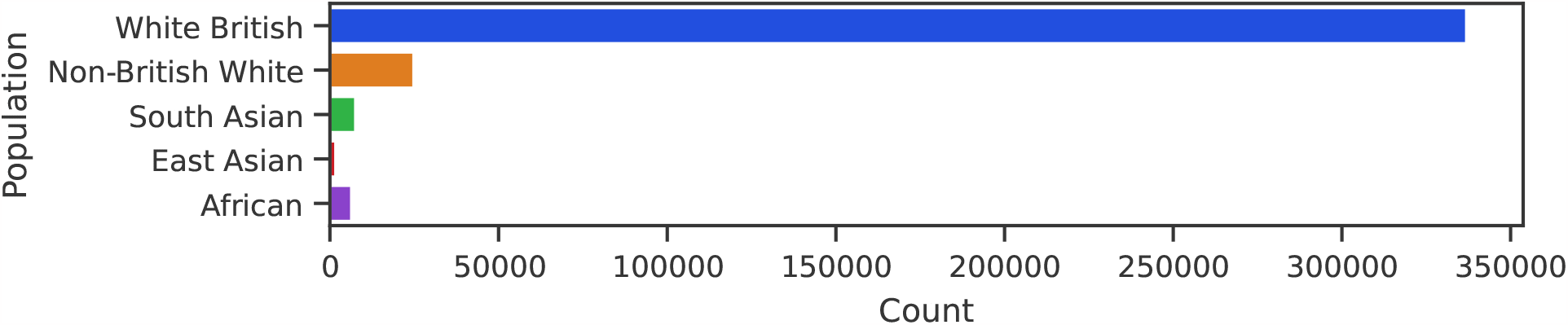
Broad population counts in the UK Biobank.^2^ Genetically inferred populations groups from the Global Biobank Engine.^4^

**Fig. 2.**
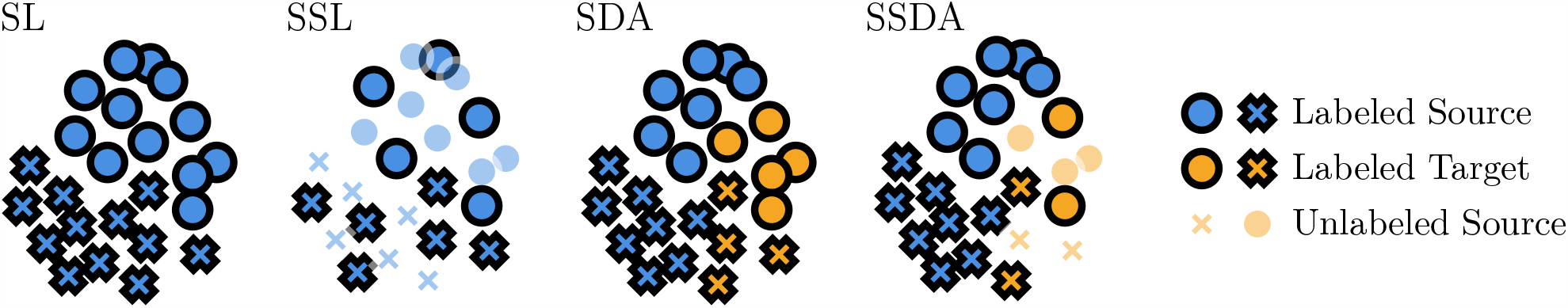
Illustration of supervised learning (SL), semi-supervised learning (SSL), supervised domain adaptation (SDA), and semi-supervised domain adaptation (SSDA) in the case of binary classification. Circle and cross markers represent negative and positive classes, opaque and transparent markers represent labeled and unlabeled points, and blue and orange markers represent source and target domains, respectively.

In this work, we propose PopGenAdapt, a semi-supervised domain adaptation (SSDA) method that can also exploit the available unlabeled data from underrepresented populations to improve the performance of phenotype prediction models. On the one hand, the semi-supervised nature of the proposed method makes possible the use of unlabeled data from underrepresented populations, as well as labeled data from large biobanks. On the other hand, the use of domain adaptation techniques makes it possible to still take advantage of the vast amount of data from individuals of European ancestry (the source domain), but to adapt the model predictions for a particular underrepresented population (the target domain). While SSDA has been previously applied to other types of data such as image and text, its application in genetics remains largely unexplored.

We adapt methods proposed for SSDA in computer vision for genotype-to-phenotype prediction and evaluate them in underrepresented population groups from Nigeria, Sri Lanka, and Hawaii. Our results predicting phenotypes including hypertension, diabetes, myxoedema, and asthma, demonstrate that SSDA can significantly enhance the performance of genotype-to-phenotype models in underrepresented populations, suggesting a promising direction for developing better machine learning models for diverse populations.

## 2. Background

### 2.1. Genotype-to-Phenotype Prediction

DNA is the hereditary material in humans and all living organisms, contributing to essential functions and appearance. While most positions in the DNA sequence are identical between individuals of the same species, some vary. Out of more than 3 billion positions, a typical human genome differs from the reference genetic sequence at 4 to 5 million sites (*∼*1.5%).^5^ In total, more than 600 million variable positions have been identified across different humans.^6^ These variable positions are called single nucleotide polymorphisms (SNPs) and can be encoded as a ternary sequence, representing the counts of non-reference variants at each position, with 0 indicating that both maternal and paternal positions match the reference genome, 1 indicating that only maternal or paternal positions match, and 2 indicating that both are alternative variants.

Phenotypes are the observable characteristics of an organism that result from the interaction between its genotype (the genetic makeup determined by its DNA sequence) and the environment. These characteristics comprise physical and behavioral traits, as well as risk of developing certain diseases. Both the frequency distribution of genomic variants, and as a result, the distribution of phenotypes, vary across different populations. As a consequence, most studies developed for a particular population do not generalize well to other population groups.^1^

The goal of genotype-to-phenotype prediction is to use the genetic variation (SNP sequences) to estimate the phenotypes of an individual. Multiple machine learning models have been applied to solve this task, either using general-purpose methods like logistic regression, gradient boosting machines, or neural networks,^7,8^ or through linear models specifically tailored to genetic data, such as PRS-CS,^9^ SBayesR,^10^ or snpnet.^11^

### 2.2. Semi-Supervised Domain Adaptation

Supervised learning is the framework most often adopted to train predictive models by using input samples and label pairs. However, in many real-world scenarios, such as in biomedical applications, obtaining labeled data can be challenging, involving time-consuming and expensive collection procedures. This limitation suggests the application of semi-supervised learning techniques, which can leverage both labeled and unlabeled data for training, providing better generalization than traditional supervised learning approaches.^12^

Both supervised and semi-supervised methods assume that the distribution of the training data (source domain) is the same as the one found during real-world deployment (target domain). However, this is not always the case, leading to distribution shifts that can drastically decrease the predictive performance. In order to address this shift, domain adaptation techniques have been proposed to properly adjust the models to bridge the gap between distributions and achieve accurate predictions in both the source and target domains.

Semi-supervised domain adaptation (SSDA) combines both semi-supervised learning and domain adaptation paradigms. The goal of SSDA is to leverage labeled data from a source domain, unlabeled data from the target domain, and a limited set of labeled data from the target domain, in order to obtain a machine learning model that achieves good performance within both domains.

In this paper, we adapt for genotype-to-phenotype prediction the state-of-the-art method of SSDA via Minimax Entropy (MME)^13^ with Source Label Adaptation (SLA),^14^ which was originally proposed in computer vision, considering different image domains, like photos, drawings, or paintings. Here, instead, we will consider different domains to be different populations.

#### 2.2.1. Minimax Entropy

Minimax Entropy (MME, Ref. 13, Fig. 3) proposes to use a neural network model consisting of a feature extractor F and a classifier *C*. At the output of F, *ℓ*_2_ normalization and temperature scaling are applied, inspired by Ref. 15. In the original work, F is a pre-trained ResNet34,^16^ an image classification network, and *C* is a single layer which takes 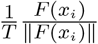as input and outputs 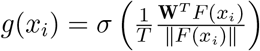.The weight vectors W = [*w*_1_,…, w_*k*_] can be regarded as a representative point of each class *k*, or “prototype”.

**Fig. 3.**
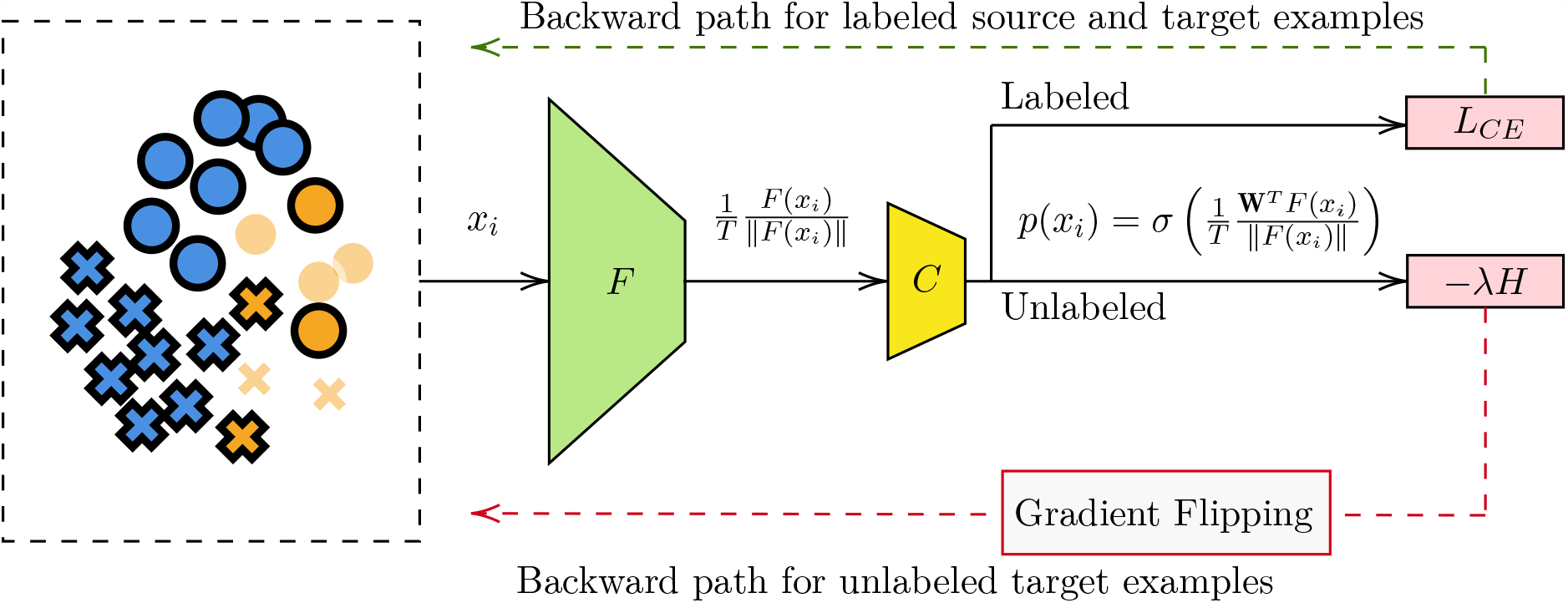
Overview of the model architecture and minimax entropy proposed in Ref. 13.

Both *C* and *F* are trained to classify labeled examples correctly by minimizing the cross-entropy loss *L*_*CE*_ on the labeled data, from both the source and target domains. However, to avoid overfitting on the source domain, which contains a larger amount of samples, as well as to take advantage of the unlabeled target data, it has been proposed to use an adversarial regularization term, the Minimax Entropy. MME is formulated as adversarial training between *F* and *C*, in which *F* is trained to minimize the conditional entropy *H* of the neural network predictions from unlabeled target data *p*(*x*_*t*_), whereas *C* is trained to maximize the entropy of the predictions *p*(*x*_*t*_). This adversarial learning forces *F* to learn discriminative features, while *C* estimates domain-invariant prototypes reducing the overfitting to the source domain. The overall adversarial learning objective functions are:

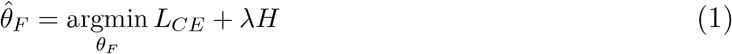

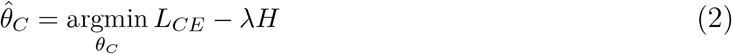

where *λ* is a hyperparameter to control the tradeoff between classification on labeled examples and the minimax entropy training. To simplify the training process, MME makes use of a gradient reversal layer^17^ to flip the gradient between *C* and *F* with respect to *H*, allowing to perform the minimax training with a single forward and backward pass.

#### 2.2.2. Source Label Adaptation

Source Label Adaptation (SLA, Ref. 14) is a framework that considers source data as a noisily-labeled version of the target data and gradually adapts the source labels to the target space. Specifically, inspired by Refs. 18,19, for each source point 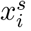, one constructs a modified source label 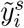 by combining, with a tradeoff ratio α, the original source label 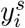 and the prediction of a source label adaptation model *p*:

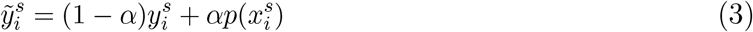

Note that *p* cannot be the current unadapted model *g* as it would overfit to the source data due to the larger number of samples, resulting in almost no effect. Thus, it has been proposed to train on the target domain data. However, to avoid simple memorization of the target data due to the low number of labeled samples available, it has been proposed to use a prototypical network (protonet),^20^ a model for few-shot learning. Given a feature extractor *F*, the prototype of class k is defined as the center of features with the same class:

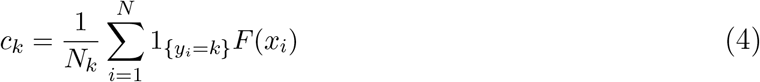

Then, a protonet produces a distribution over classes for a query point *x*_*i*_ based on a softmax with temperature *τ* over the Euclidean distances to the prototypes in the embedding space:

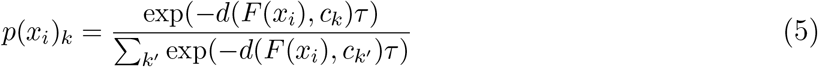

Moreover, in Ref. 14 it is proposed to derive the prototypes using the unlabeled data available by using, for each unlabeled target instance 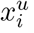, pseudo labels 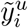 computed by the current model g:

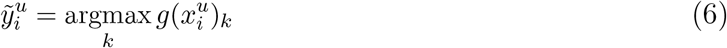

Using these pseudo labels, we can get pseudo centers by Eq. 4, and further define with them a Protonet with Pseudo Centers (PPC) by Eq. 5. Next, the PPC is applied to Eq. 3 to compute the modified source labels 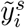 for each source instance 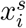. Finally, the real source labels 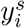 are replaced by the cleaned source labels 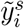 in the computation of the cross-entropy for the labeled source part of the whole dataset. The loss for labeled target data can still be a standard cross-entropy loss. Other loss terms can still be included, like the minimax entropy proposed in Ref. 13.

In practice, the SLA framework is only applied after W warmup steps in which the model is trained normally with the original source labels to obtain an initial robust model, and then the pseudo labels are only recomputed every I steps for efficiency. Since the SLA^14^ paradigm of considering the source labels as noisy from the target domain viewpoint and cleaning them is orthogonal to the ideas in MME,^13^ both approaches can be combined to get superior results. We refer to this combination as MME-SLA.

### 2.3. Semi-Supervised Learning and Domain Adaptation for Genotype-to-Phenotype Prediction

Semi-supervised learning techniques have been previously applied in genotype-to-phenotype prediction. For example, Ref. 21 proposed a method to predict the residual feed intake in dairy cattle using both labeled and unlabeled samples. However, the samples are assumed to be from the same domain, so the method would still have the problem of not generalizing to other populations.

Likewise, domain adaptation techniques have also been applied in genotype-to-phenotype prediction. For instance, Refs. 22–24, proposed several transfer learning techniques to also improve prediction performance for underrepresented populations. However, the proposed approaches cannot utilize unlabeled samples, thereby still grappling with the scarcity of labeled data from underrepresented populations. Consequently, the achieved performance improvement remains limited. To our knowledge, this is the first work to combine both approaches by applying semi-supervised domain adaptation for genotype-to-phenotype prediction.

## 3. Method

### 3.1. Data

We apply these methods to predict multiple disease outcomes, including hypertension, diabetes, myxoedema, and asthma, for individuals from populations underrepresented in commonly used datasets, including Nigeria, Sri Lanka, and Hawaii, available in the UK Biobank^2^ and the PAGE study.^25^ In order to have meaningful results, we limit the phenotypes to these four, as they have a high enough case count within the three target populations. For each phenotype, we use balanced data from white British individuals as the source domain, obtained by removing samples from the majority class. Then, for each phenotype and target population, we use the labeled source domain data as well as labeled and unlabeled data from the target domain. To test the method’s efficacy, we use a subset of labeled data exclusive to the target underrepresented population.

To establish which samples constitute the target population domain, we propose two approaches, which show two different ways in which we can take advantage of the availability of datasets, even when the labeled data in the target domain is very scarce. The first approach is adopted for the Nigerian and Sri Lankan populations, and the second one for the Hawaiian population.

The first approach only uses data from the UK Biobank.^2^ To establish which samples constitute the target population, we combine the genetically inferred ancestry available from the Global Biobank Engine^4^ (white British, non-British white, South Asian, East Asian, or African) and the country of birth reported in the UK Biobank.^2^ We use both fields because the inferred genetic ancestry provides a continental-level description, encompassing many regions within each label. On the other hand, the country of birth alone is not representative of the ancestry composition within the UK Biobank due to high selection bias, as the samples were collected in assessment centers in the United Kingdom, so many individuals in the data born outside the United Kingdom are still of English genetic ancestry. By filtering both by inferred population group and country of birth, we ensure that the definition of the target domain is precise.

In particular, for the case of Nigeria, we only keep the samples that are of African genetic ancestry and born in Nigeria, and for the case of Sri Lanka, the samples that are of South Asian genetic ancestry and born in Sri Lanka. This results in a total of 852 samples for the Nigeria group and 535 samples for the Sri Lanka group. Once we have the samples from the target domain, since the UK Biobank has phenotype labels for all the samples, we artificially unlabel half of them for the purpose of evaluating the proposed method. For training, we use all the unlabeled samples plus only 10 labeled samples from the target domain, 5 negative and 5 positive, alongside all the labeled samples from the source domain. The rest of the labeled individuals of the target domain are split into two equal parts using stratified sampling to create the validation and test sets. Note that we can only use labeled data for validation and testing.

The second approach to define the target domain shows how additional unlabeled datasets can be employed. To achieve this, in addition to the UK Biobank,^2^ we use a dataset of SNP sequences (without phenotype labels) of 5,862 Native Hawaiian individuals from the PAGE study.^25^ In this setting, we only have unlabeled data from the target population. Note that we cannot use the country of birth field, as the people born in Hawaii are labeled as born in the USA. To have labeled data in the target domain, we propose to use the nearest neighbor of each sample from the Hawaiian dataset within the UK Biobank, excluding the white British individuals to avoid having repeated samples in both the source and target domains. For efficiency, we compute the distances between samples on the first 50 principal components, instead of using the raw SNP sequences. After removing duplicated individuals that are the nearest neighbor to more than one sample from the Hawaiian dataset, we obtained 1,689 labeled samples. While it is unlikely that the UK Biobank contains this many individuals of Hawaiian ancestry, the closer distribution of these samples to the Native Hawaiian population makes the domain more apt to model them than using samples of predominantly European ancestry.

The second approach to defining the target domain is less accurate than the first one, as it includes samples from other similar populations. However, it has the advantage that it results in a larger number of samples, which can be helpful for unbalanced phenotypes with a low positive case count, and to counteract the effect of having a noisier target domain definition. In this scenario, we use 50% of the labeled target samples for the training set, 25% for the validation set, and 25% for the test set. Note that the unlabeled samples used for training are the ones from the Hawaiian dataset from the PAGE study.^25^

We use the variants that are both in the UK Biobank data and the Hawaiian dataset, resulting in 83,362 overlapping SNPs. Note that we decided to not impute the SNPs outside the intersection to avoid introducing a bias. Most algorithms to perform statistical imputation are based on the available samples, and since in this scenario most of them are not from the underrepresented population, the imputation could result in incorrect values.

### 3.2. Model

We adopt the MME-SLA^13,14^ method originally proposed for classification tasks in computer vision for genotype-to-phenotype prediction by replacing the ResNet34^16^ backbone model used in the original works with a multi-layer perceptron (MLP). Specifically, we use a 4-layer MLP with GELU activations,^26^ layer normalization,^27^ and a residual connection^16^ between the output of the first layer and the input of the last one. The choice of activation and the use of layer normalization and a residual connection is commonly adopted in modern architectures such as Transformers^28^ and has been proven to help improve the performance of the models, as well as their stability during training. The initial layer of the network takes an input size corresponding to the number of SNPs and reduces it to a hidden size of 256. Subsequently, the two middle layers maintain the same input and output dimensions of 256. Next, before the last layer, *ℓ*_2_ normalization and temperature scaling with T = 0.05 is applied, as proposed in Refs. 13,15. Lastly, the last layer, which acts as the classifier, produces an output size equivalent to the number of classes. We call the complete model PopGenAdapt.

The backbone MLP model without the MME-SLA components is also used as the baseline model to compare how applying SSDA improves against a typical supervised learning approach.

We train the baseline and PopGenAdapt models for each combination of target population and phenotype using a batch size of 64, the AdamW optimizer^29^ with weight decay of 0.01, and the same learning rate scheduler used in Ref. 14. We use randomized hyperparameter search to tune several hyperparameters. For the baseline method, we only tune the learning rate. For PopGenAdapt, we tune both the learning rate and the MME-SLA^13,14^ hyperparameters (*λ, α, τ, W*, and *I*). Table 3 shows the hyperparameter space from which the values were sampled. Our experiments showed that the resulting performance is highly sensitive to the choice of hyperparameters, as also pointed out in the paper which introduced SLA.^14^ We select the model with the best validation AUROC on the target domain and perform the final testing on a separate hold-out test set also using AUROC on the target domain.

**Table 1.** Size of sets used for training and evaluation for each population. Note that a combination of white British as the source domain plus another population as the target domain is always used.

| Population | Training labeled | Training unlabeled | Val. + Test labeled | Total |
| --- | --- | --- | --- | --- |
| White British | * | 0 | 0 | * |
| Nigeria | 10 | 213 | 106 + 107 | 852 |
| Sri Lanka | 10 | 134 | 67 + 67 | 535 |
| Hawaii | 822 | 5,862 | 412 + 413 | 7,507 |

\*White British set size depends on class balancing, but in all cases is >40,000.

**Table 2.** Case counts for each disease and population (only considering labeled samples).

| Population | Hypertension | Myxoedema | Diabetes | Asthma |
| --- | --- | --- | --- | --- |
| White British* | 114,687 (50.00%) | 21,471 (50.00%) | 23,099 (50.00%) | 45,192 (50.00%) |
| Nigeria | 105 (47.08%) | 11 (4.93%) | 38 (17.04%) | 21 (9.41%) |
| Sri Lanka | 60 (41.66%) | 18 (12.50%) | 41 (28.47%) | 29 (20.13%) |
| Hawaii | 535 (31.68%) | 69 (4.09%) | 153 (9.05%) | 197 (11.66%) |

\*White British phenotypes are balanced by undersampling the negative class.

**Table 3.** Definition of the distribution of the hyperparameter space.

| Hyperparameter | Probability distribution |
| --- | --- |
| Learning rate | LogUniform($10^{-5}$, $10^{-2}$) |
| MME tradeoff $\lambda$ | Uniform(0, 1) |
| SLA mix ratio $\alpha$ | Uniform(0, 1) |
| SLA temperature $\tau$ | Uniform(0, 1) |
| SLA warmup $W$ | UniformChoice({100, 500, 1000, 2000, 5000}) |
| SLA update interval $I$ | UniformChoice({5, 10, 100, 500, 1000, 2000, 5000}) |

Note that while PopGenAdapt employs both the labeled and unlabeled samples, the baselines are trained on the subset that is labeled, as it has no way of using the unlabeled samples.

The training and inference was performed with an NVIDIA GeForce GTX 1080 Ti GPU (11 GB), and took between 10 and 50 minutes, depending on the number of samples and the hyperparameters, for each configuration.

## 4. Results

We compare PopGenAdapt with the baseline model consisting only of the backbone MLP (MLP Base), as well as with the state-of-the-art genotype-to-phenotype snpnet^11^ model, and PRS-CSx,^30^ which is an extension of PRS-CS^9^ to improve polygenic prediction in ancestrally diverse populations. Note that since snpnet and PRS-CSx are supervised models, like in the case of the baseline model, they can not exploit the unlabeled samples.

We show the results obtained for each of the four phenotypes on the three tested target underrepresented populations in Tables 4–6.

**Table 4.** AUROC for the Nigerian population.

| Method | Hypertension | Myxoedema | Diabetes | Asthma | Average |
| --- | --- | --- | --- | --- | --- |
| snpnet<sup>11</sup> | 0.4647 | 0.7949 | 0.4699 | 0.4646 | 0.5485 |
| PRS-CSx<sup>30</sup> | 0.3884 | 0.1827 | **0.6046** | 0.4306 | 0.4016 |
| MLP Base | **0.5488** | 0.4423 | 0.5376 | 0.4886 | 0.5043 |
| PopGenAdapt | 0.5088 | **0.8109** | 0.5516 | **0.5619** | **0.6083** |

**Table 5.** AUROC for the Sri Lankan population.

| Method | Hypertension | Myxoedema | Diabetes | Asthma | Average |
| --- | --- | --- | --- | --- | --- |
| snpnet<sup>11</sup> | 0.4500 | 0.4631 | 0.4603 | 0.5871 | 0.4901 |
| PRS-CSx<sup>30</sup> | 0.4852 | 0.4863 | 0.3991 | 0.5379 | 0.4771 |
| MLP Base | **0.5898** | 0.5137 | 0.5952 | 0.5939 | 0.5731 |
| PopGenAdapt | 0.5778 | **0.5902** | **0.6723** | **0.6091** | **0.6123** |

**Table 6.** AUROC for the Hawaiian population.

| Method | Hypertension | Myxoedema | Diabetes | Asthma | Average |
| --- | --- | --- | --- | --- | --- |
| snpnet<sup>11</sup> | 0.6132 | **0.6148** | 0.5235 | 0.4857 | 0.5593 |
| PRS-CSx<sup>30</sup> | 0.5423 | 0.5881 | 0.4577 | 0.5087 | 0.5242 |
| MLP Base | 0.6104 | 0.5162 | 0.5041 | 0.5666 | 0.5493 |
| PopGenAdapt | **0.6135** | 0.5556 | **0.5791** | **0.5811** | **0.5823** |

PopGenAdapt outperforms snpnet, PRS-CSx, and the baseline model on average and in the majority of evaluated scenarios. Moreover, we observe that snpnet, PRS-CSx, and the baseline model obtain in multiple cases an AUROC below 0.5, indicating a predictive performance worse than the one obtained by random guessing. We note that this does not happen in any of the experimented cases for PopGenAdapt. Considering that snpnet and the MLP baseline methods do not perform any type of domain adaptation, it makes sense for this to happen, as the models are tested on a domain that differs from the one in which most of the training samples are.

We hypothesize that a possible reason for the poor performance of snpnet on non-European populations is due to the use of the lasso in the method, which performs SNP selection, thus excluding completely some variants. Possibly, as the training data is mostly from the source domain, many of the SNPs excluded are the ones that remain useful for making the prediction on the target population. All this reflects the usefulness and need for domain adaptation techniques to be used when the data of the target domain is limited, like in the case of underrepresented populations.

Furthermore, PRS-CSx has poor performance in most settings. We believe that the small number of samples of the target population still had an effect on this case, reflecting the value of incorporating unlabeled samples when the labeled data is scarce. Another possible limitation that could result in bad performance is the use of a relatively small number of SNPs, although this is shared across all four methods.

Finally, we also observe that the supervised methods suffer less in the Hawaiian dataset, probably due to the higher number of labeled samples for training that are used in this scenario and the fact that the Hawaiian target domain is less precise, resulting in less advantage for domain adaptation. The target domain, in this case, is less precise due to the nearest neighbor approach used to establish the labeled samples, as well as due to the fact that Pacific Islanders are admixed populations, resulting in more variability across the samples, as can be observed in Fig. 4.

**Fig. 4.**
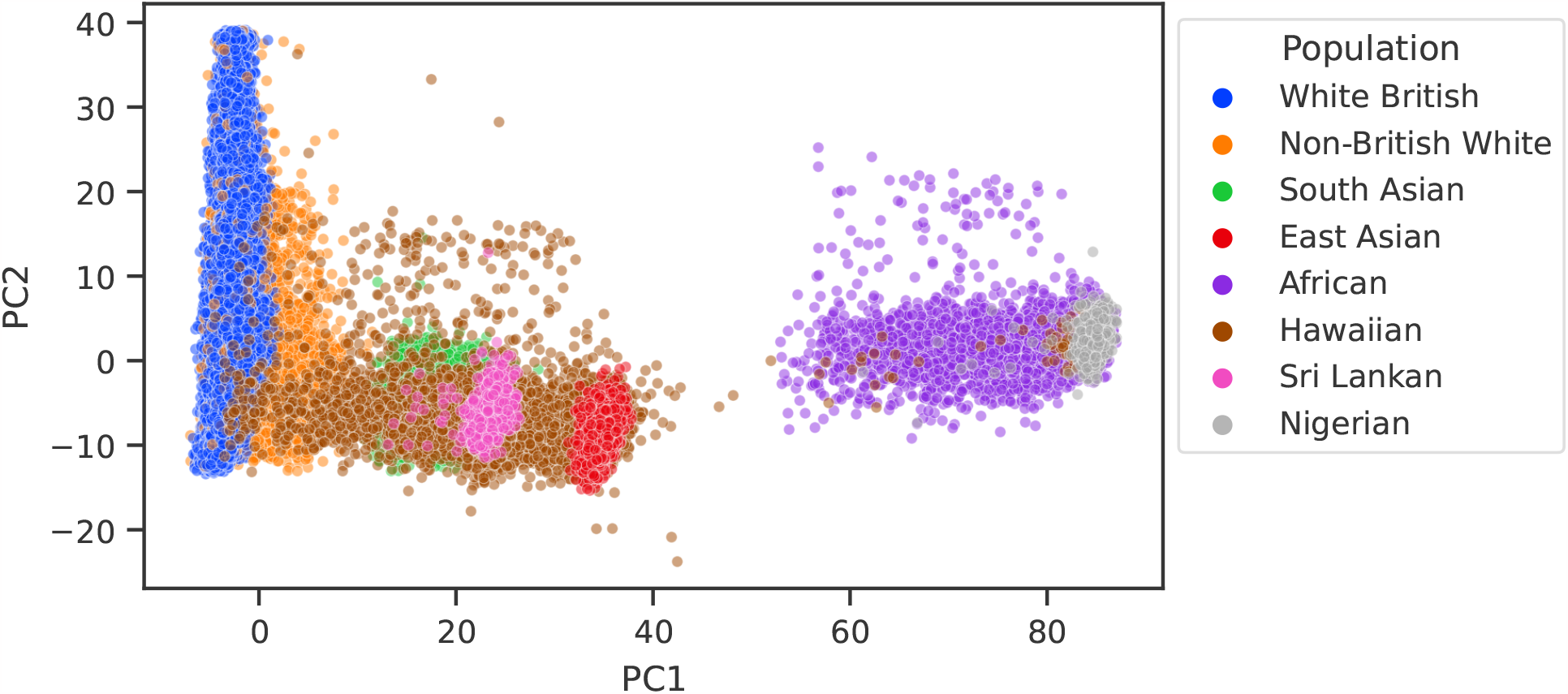
Two-dimensional PCA projection of the samples in the UK Biobank and the Hawaiian dataset. The PCA was fitted with only the samples from the UK Biobank. Note that all samples marked as Sri Lankan fall within the South Asian genetic ancestry cluster, and all the Nigerian ones fall within the African cluster.

**Fig. 5.**
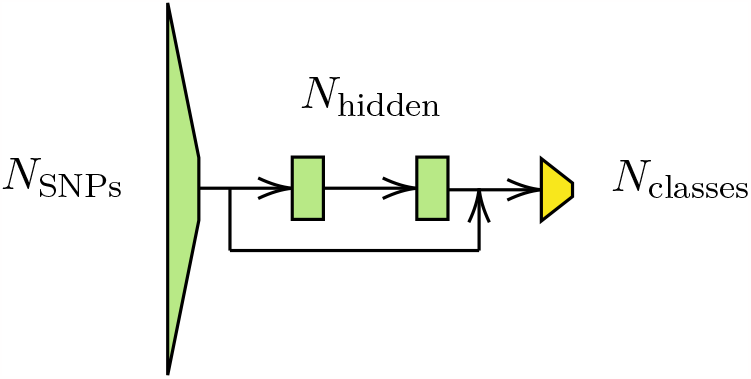
Diagram of the backbone MLP model for PopGenAdapt.

## 5. Conclusion

In this work, we presented PopGenAdapt, a model that applies semi-supervised domain adaptation techniques for genotype-to-phenotype prediction. We also proposed two approaches to set the target domain samples and evaluated the model to predict several disease outcomes in three different underrepresented populations. The results show that by using SSDA on underrepresented populations, the prediction performance can be improved over state-of-the-art supervised methods. Consequently, we show SSDA is a promising technique to help overcome health disparities in precision medicine by exploiting the availability of unlabeled data from underrepresented populations while still taking advantage of the greater magnitude of labeled data available from populations of European ancestry.

Nonetheless, there are still some limitations and avenues for future work. Due to the limited data on the underrepresented population we had available from the UK Biobank,^2^ we did not study the influence the ratio of labeled and unlabeled samples could have on the attained performance, as using more samples for training would have left too few for validation and testing. Moreover, the scalability of the method to a larger number of SNPs also remains to be assessed. Further work on the approach could also include the possibility of learning from GWAS summary statistics instead of the SNP sequences or to also support continuous phenotypes apart from categorical ones. Possibly, there is also room for improvement on the base model used, as more powerful deep learning architectures could be evaluated. Furthermore, considering the integration of PopGenAdapt on emerging paradigms such as federated learning or differential privacy^31^ could further enhance the applicability of the method in biomedical research and healthcare.

## Acknowledgments

We would like to thank Mark A. Penjueli for data pre-processing of the PAGE study dataset. The computing for this project was performed on the Sherlock cluster from Stanford University and on the TSC CALCULA cluster from the Department of Signal Theory and Communications (TSC) of the Polytechnic University of Catalonia (UPC). We would like to thank both institutions for supplying computational resources and support that contributed to these results. This work was partially supported by NIH under award R01HG010140 and by a grant from the Stanford Institute for Human-Centered Artificial Intelligence (HAI). This research has been conducted using the UK Biobank Resource under Application Number 24983.

